# A spatial cell culture model for predicting chemotherapy dosing strategies

**DOI:** 10.1101/561746

**Authors:** Shu Zhu, Dhruba Deb, Tal Danino

**Author notes:** Correspondence should be addressed to T.D.

## Abstract

Predicting patient responses to chemotherapy regimens is a major challenge in cancer treatment. To do this requires quantitative mathematical models to predict optimal dose and frequency for a particular drug, and experimental model systems such as three-dimensional organoids that accurately recapitulate the tumor microenvironment and heterogeneity. However, tracking the spatial dynamics of multiple cell types in three-dimensions can be a significant challenge in terms of time and throughput. Here we develop a two-dimensional system that allows for simple tracking of cell populations via fluorescence microscopy for modeling spatial dynamics in tumors. We first develop multiple 4T1 breast cancer cell lines resistant to varying concentrations of doxorubicin, and demonstrate how well mixed and spatially heterogeneous populations expand in a two-dimensional colony. We subject cell populations to varied dose and frequency of chemotherapy and measure colony growth radius and populations. We then build a mathematical model to describe the dynamics of both chemosensitive and chemoresistant populations, where we determine which number of doses can produce the smallest tumor size based on parameters in the system. In the future, this system can be adapted to quickly optimize dosing strategies in the setting of heterogeneous cell types or patient derived cells with varied chemoresistance.

## Introduction

Chemotherapy dosing strategies have primarily followed a maximum tolerated dosage (MTD) approach, in which patients are given high doses of chemotherapy to kill as many tumor cells as possible (1–5). This approach is based on early assumptions that tumors are composed of a homogenous, exponentially growing cell population and thus maximum doses are likely to cause the highest disease eradication (6, 7). However, tumors are composed of genetically heterogeneous cells with varied chemoresistance, and further diversity in chemoresistance can arise due to drug selection pressure from traditional high dose treatments (8–10). Thus, while eradicating chemosensitive cells, the MTD approach can lead to faster growth of chemoresistant cell populations, leading to faster relapse and eventually worse outcomes (11). Furthermore, due to the severity of side-effects at high doses, administration of chemotherapy using the MTD approach is typically separated by timescales as large as weeks to allow normal tissue to recover (12). In contrast to MTD, metronomic chemotherapy (MCT) (13, 14), where chemotherapy is delivered at lower doses more frequently, has been explored and demonstrated in some cases to be a more optimal dosing strategy in terms of both measures of efficacy and resistance development (15–18).

Since chemotherapeutic dosing schedules are often determined empirically in clinical trials, the discovery of optimal dosing strategies such as MCT for individual drugs has been limited. Several *in silico* studies coupled with *in vitro* assays have demonstrated that intra-tumor heterogeneity and dosing strategy can affect tumor response to chemotherapy (9, 15, 19, 20). However, *in vitro* drug efficacy assays are typically performed on homogenous, chemosensitive tumor cells, which fail to incorporate tumor spatial heter ogeneity and thus are not ideal for investigating the spatial competition and dynamic interaction between drug sensitive and resistant cell subpopulations in response to chemotherapy. Inevitably these models are not predictive of drug efficacy or dosing schedules, particularly in patients who have developed drug resistance from previous treatments (6, 21). In order to recapitulate the complexity of the tumor microenvironment, three-dimensional (3D) multicellular spheroid and organoid models have been employed to incorporate transport dynamics and intricate cell-cell interactions that are naturally presented *in vivo* (20, 22–24). However, it is challenging to incorporate clonal heterogeneity in a spatially-controlled manner or visually track resistant clone trajectories in 3D cell culture models.

Here we present a novel 2D culture system that allows for tracking dynamics of subclonal populations in a growing colony. This model incorporates cell-cell interactions and clonal chemoresistance heterogeneity, where colony growth rate can be simply measured as the change in colony diameter without going through tedious cell counting process. Using this system, it is possible to control the initial resistant cell proportion, as well as the spatial distribution during the seeding process to recapitulate various tumor spatial structures. The spatial evolution of resistant clone and interaction between chemoresistant and chemosensitive clones can be easily visualized using standard epifluorescence microscopy. The colony expansion mimics tumor expansion for at least 3 weeks in standard tissue culture wells, which makes long-term analysis of chemotherapy dosing schedules possible. Finally, we utilize the behavior of this 2D culture system to build a mathematical model and study the effect of various chemotherapy dose and schedules on the tumor growth.

## Materials and methods

### Development of chemoresistant breast cancer cell lines

Mice triple negative breast cancer cell line 4T1 was obtained from ATCC and was cultured in RPMI cell culture medium (RPMI 1640, Thermo Fisher, MA) supplemented with 10% fetal bovine serum (Thermo Fisher, MA) and penicillin-streptomycin (100 IU/ml). Cultured cells were maintained in a controlled humidified atmosphere of 5% CO_2_ in air at 37°C. Cells were subcultured every 3-4 days when reached over 80% confluency. Drug sensitivity of wild type cells were evaluated using a colorimetric cell proliferation assay (MTT assay, V13154, Thermo Fisher, MA) and a dose response curve was constructed by culturing cells and evaluating cell viability in the presence of various chemotherapy drug concentrations. In order to establish chemoresistant cell lines, wild type cells were initially cultured in the presence of a very low concentration of chemotherapy (approximately 5% of IC_50_). Cell medium was switched every 2-3 days until 75-80% confluency was reached. Cells were subsequently disassociated and subcultured and the viability of collected cells was evaluated using trypan blue staining. Drug concentration in the subcultured cells was doubled when over 90% of collected cells were viable. Otherwise, cells were subcultured and maintained at the same concentration for additional 1-2 passages until desired viability was achieved. Cell drug sensitivities were evaluated at each passage using MTT assay and cells with selected drug resistance were cryopreserved. In the present work, the resistance level of cells is defined as the highest chemotherapy concentration the cells were exposed to while being cultured. For example, the label 4T1-Dox [50 nM] means 4T1 cells that are able to replicate and grow in the presence of 50 nM doxorubicin (Sigma-Aldrich, MO) and the viability of cells at confluency are over 90%.

### 2D radial growth cell culture model

Cryopreserved cells with desired chemoresistance were thawed and cultured in the presence of chemotherapy with matched concentration for at least 2 passages prior to 2D radial growth experiments for cells to reach stable growth state. Harvested cells were then resuspended in cell culture medium with a density of 10^6^ cells/mL. Single drops of 20 μL cell suspension were subsequently seeded in the center of each well on a surface treated 12-well plate. Significant spreading of the drop was not observed due to the hydrophobicity of treated well plate surface. Cells were allowed to settle for 24 hours under normal culture condition (5% CO2 in humidified air, 37 °C) followed by a gentle PBS wash to remove dead or loosely attached cells. Fresh medium was then added to cover the whole well. At this point, a single colony of cells should have already appeared in the center of each well, the diameter of the colony is defined as the initial diameter and can be adjusted by varying the combination of seeding drop volume and seeding cell density. The seeded colonies would start expanding radially once the plates were returned to standard culture condition. Gentle PBS wash and refill of fresh medium was done every 24 hrs (Fig 1, case I).

**Fig 1.**
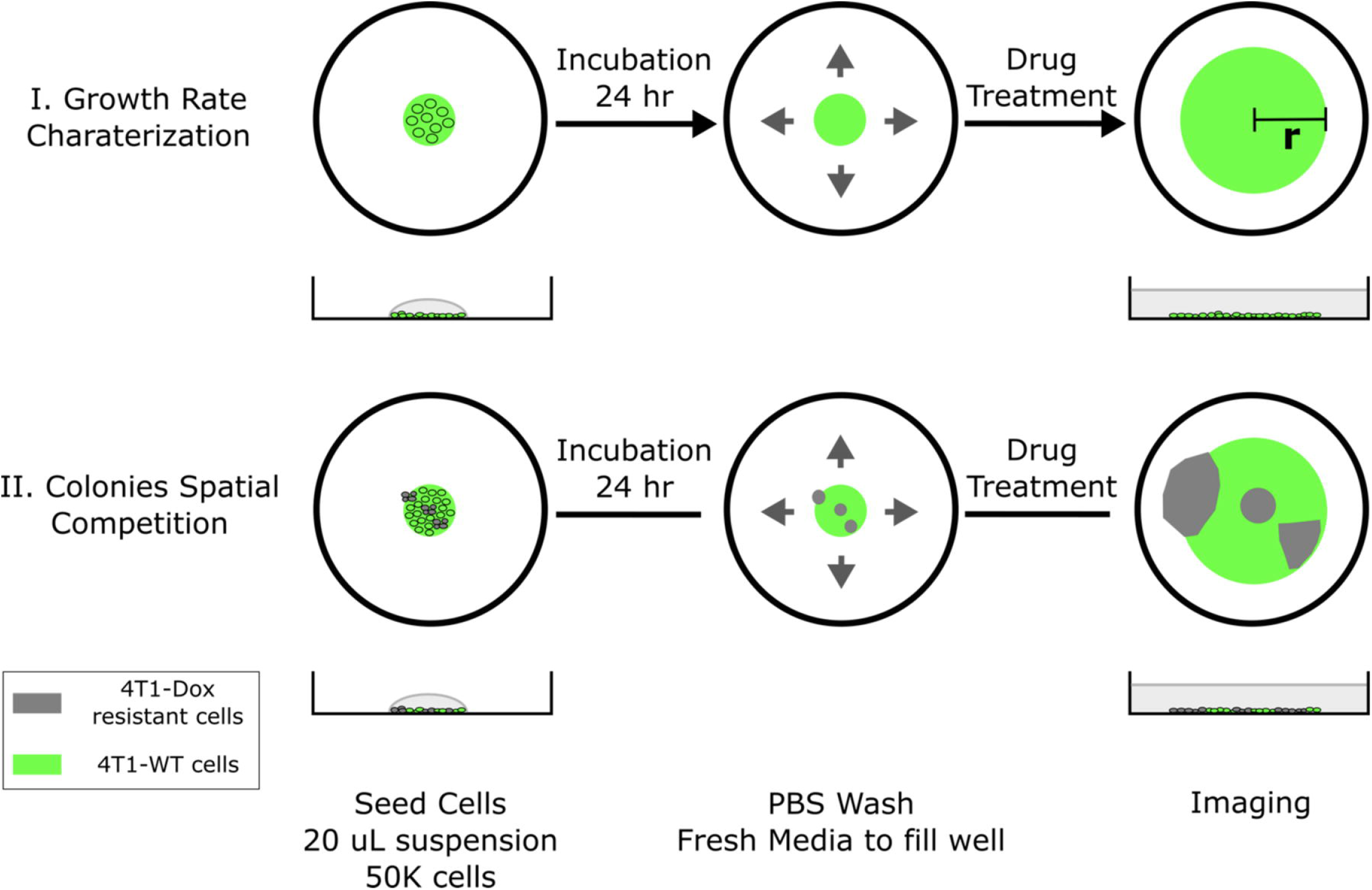
Schematic of 2D spatial colony system. (top) Chemosensitive 4T1 cells are seeded in a well plate and grow radially, where colony radius and cell fluorescence are tracked over time. (bottom) A mixture of chemosensitive and chemoresistant cells are premixed or spotted sequentially to establish a spatial competition model between cell populations.

The cell seeding procedure can be modified to incorporate spatial heterogeneity in drug resistance across the colony. Single drops of 20 μL wild type 4T1 cell suspension (10^6^ cells/mL) were first seeded in the center of each well. Using a 10uL pipette tip, a 5 μL drop of resistant cell suspension was gently added into the center of each seeded big drop without disturbing the big drop. The tip was inserted into the big drop vertically and the resistant cell suspension was slowly injected right above the bottom of the well. Cells were allowed to settle for 24 hours before a PBS wash. Fresh medium was subsequently added to fill the whole well to initiate colony expansion. The tip can be inserted into the center or near the edge of the drop and the resulted resistant subcolony will be observed at the corresponding location within the big colony (Fig 1, case II). The volume of added resistant cell suspension defines the size of resistant subcolony. It is also possible to incorporate multiple resistant clones by adding more drops of resistant cell suspension. In the case of coculturing chemosensitive and chemoresistant cells, 4T1-citrine (fluorescent wild type 4T1 cells) were used to differentiate different cell populations.

### MTT assay

Cell viability assays were done according to manufacturer’s instructions (V13154, Thermo Fisher, MA). Briefly, cells were seeded on a 96-well plate with density of 6000 cells/well and the plate was incubated under normal cell culture condition for 24 hr before adding 10 μL of MTT reagent into each well. The well plate was immediately returned to cell culture incubator and was incubated for another 3-4 hours until visible purple precipitations show up at the bottom. Another 100 μL of detergent reagent was then added to dissolve the purple precipitations. OD measurement of the well plate was done at 570 nM wavelength using a Tecan microplate reader (Thermo Fisher, MA) and the background for each well was measured at 690 nM wavelength and was subtracted from the readings. Wells with wild-type cells were controls and each condition was tested in at least triplicate wells.

### Image Analysis

Each well was imaged using an EVOS FL Auto 2 Cell Imaging System (Thermo Fisher, MA) at desired time points. The scope and accessories were programmed using the Celleste Imaging Analysis software (Thermo Fisher, MA). Customized MATLAB code was used stitch subplot and generate images of each entire colony. Images were analyzed using Image J (NIH). Outlines of each colony were traced manually in ImageJ. Area measurements were used to calculate colony diameter assuming each colony is a perfect circle.

## Results

### Characterization of doxorubicin-resistant 4T1 breast cancer cell lines shows their presence in the 2D co-culture system can alter the overall response to doxorubicin treatment

MTT analysis confirmed that gradually increasing chemotherapy concentration in cell culture leads to the generation of 4T1 chemoresistant cell lines with different level of chemoresistance (Fig 2A). While the IC_50_ of wild type 4T1 cells was about 2 μM, the two representative resistant cell lines we created were able to maintain viability at much higher drug concentrations. For 4T1-Dox (3 μM), 4T1 cells that were cultured in the presence of 3 μM of Dox, we observed minimal reduction in cell viability even at 100 μM, which was the highest concentration tested due to the limited drug solubility.

**Fig 2.**
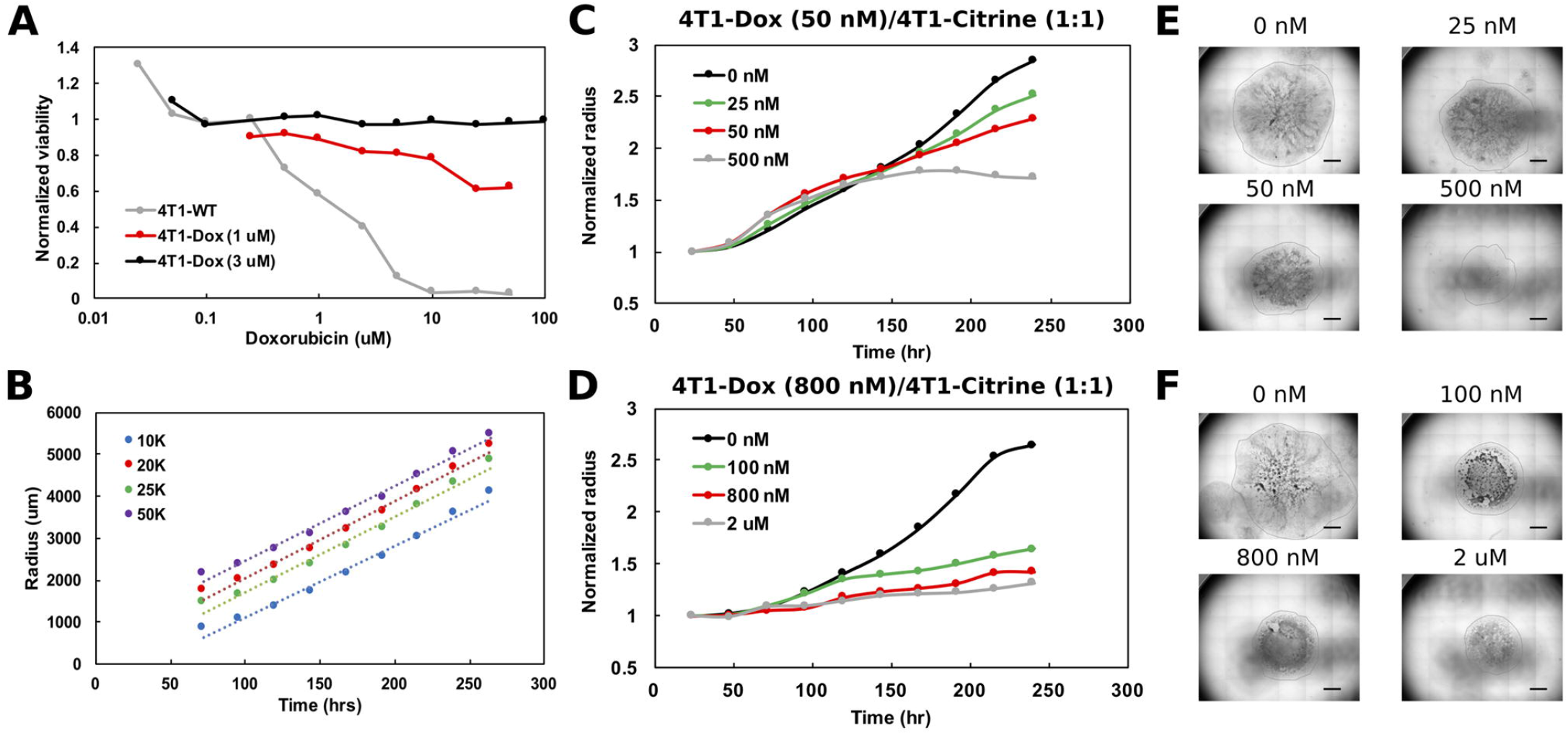
Generation and characterization of doxorubicin-resistant 4T1 cells in a 2D model system. **(A)** Viability of chemosensitive or Dox-resistant 4T1 cells across varied Doxorubicin added to culture determined by MTT assay. **(B)** Radial growth of chemosensitive colonies (4T1-WT cells) as a function of time for various seeding densities. **(C)** Colony radius normalized to starting radius in a 1:1 seeded mixture of sensitive and 50nM Dox resistant cells in the presence of 0 nM, 25 nM, 50 nM or 500 nM doxorubicin. **(D)** Colony radius normalized to starting radius in a 1:1 seeded mixture of sensitive and 800nM Dox resistant cells in the presence of 0 nM, 100 nM, 800 nM or 2 μM doxorubicin. **(E)** and **(F)** Stitched 4X brightfield images demonstrating size of colony and morphology across various seeding and chemotherapy conditions in (C) and (D), respectively. Scale bar = 1000 μm.

We next assessed the effect of initial cell density on the growth of the colonies over time. Specifically, maintaining the seeded drop volume constant at 20 μL, increasing seeded cell number (all sensitive cells) from 10K to 50K resulted in an over 2x increase in initial
colony diameter (Fig 2B). However, the colony expansion rates indicated by the change in colony diameter within a fixed time period (slope of the growth curves) were similar despite the difference in initial colony diameter. Reduction in serum level impeded colony expansion, however, where a 10x reduction in serum level was required to achieve a significant reduction in colony growth rate (Fig S1).

During later cycles of chemotherapy in patients, intratumor heterogeneity is common and sensitive and resistant cells often co-exist (25). Hence, we hypothesized that our model system could be used to test the overall tumor response to chemotherapy. Here, we seeded single drops of 20 μL wild type 4T1 cell suspension (10^6^ cells/m L) in the center of each well (~20K cells), and a varied number of resistant cells were added. Mixing resistant and sensitive cells and seeding mixed cells lead to the formation of colonies with both cell populations and resistant cells were evenly scattered across the entire colony. To mimic the clinically relevant level and range of resistance (26), we chose to utilize cells with lower resistance levels (<1 μM) for our characterization. For colonies with cells that were less resistant (4T1-Dox [50 nM], which grow at 50 nM doxorubicin), increasing the initial proportion of resistant cells had minimal impact on colony expansion in the absence of any doxorubicin treatment (Fig 2C, Fig S2 A-C). Interestingly, for colonies with cells that were more resistant to doxorubicin (4T1-Dox [800 nM], which grow at 800 nM doxorubicin), a marked reduction in colony expansion was observed when the majority of seeded cells were resistant even in the absence of drug treatment (Fig S2 F). For all tested conditions, doxorubicin treatment had minimal impact on colony expansion during the initial stage while the reduction in colony expansion was more pronounced during later time points (Fig 2C, D, Fig S2). Taken together, our data suggests that presence of doxorubicin resistant 4T1 cells along with sensitive cells can alter the overall response to doxorubicin in our model and this response varies based on the dose and time of the treatment.

### 2D co-culture system can be optimized to identify a critical drug concentration and to screen for various treatment regimens

In order to differentiate the response of sensitive and resistant cells, DAPI staining of each colony was carried out at the final time point. In the end, the whole cell population stains with DAPI while only the sensitive cells show the GFP signal. Hence, cells that were DAPI-positive, but GFP-negative were identified as the resistant cells. By measuring the occupied area of each cell population, we found high drug doses caused more reduction in final size of the colony but at a price of increased proportion of resistant cells in the colony (Fig 3). Without any drug treatment, colony expansion was relatively faster but majority of the colony was occupied by sensitive cells indicating a growth advantage of sensitive cells in the absence of drugs. The drug concentration that led to a colony composition of equal amount of chemoresistant and chemosensitive cells can be considered as a “critical concentration.” Both the initial proportion of resistant cells as well as their level of resistance determined this critical concentration and it was much lower when majority of seeded cells were chemoresistant. Interestingly, the size of the colonies in presence of highly resistant cells, 4T1-Dox [800 nM], did not reduce markedly beyond the critical concentration.

**Fig 3.**
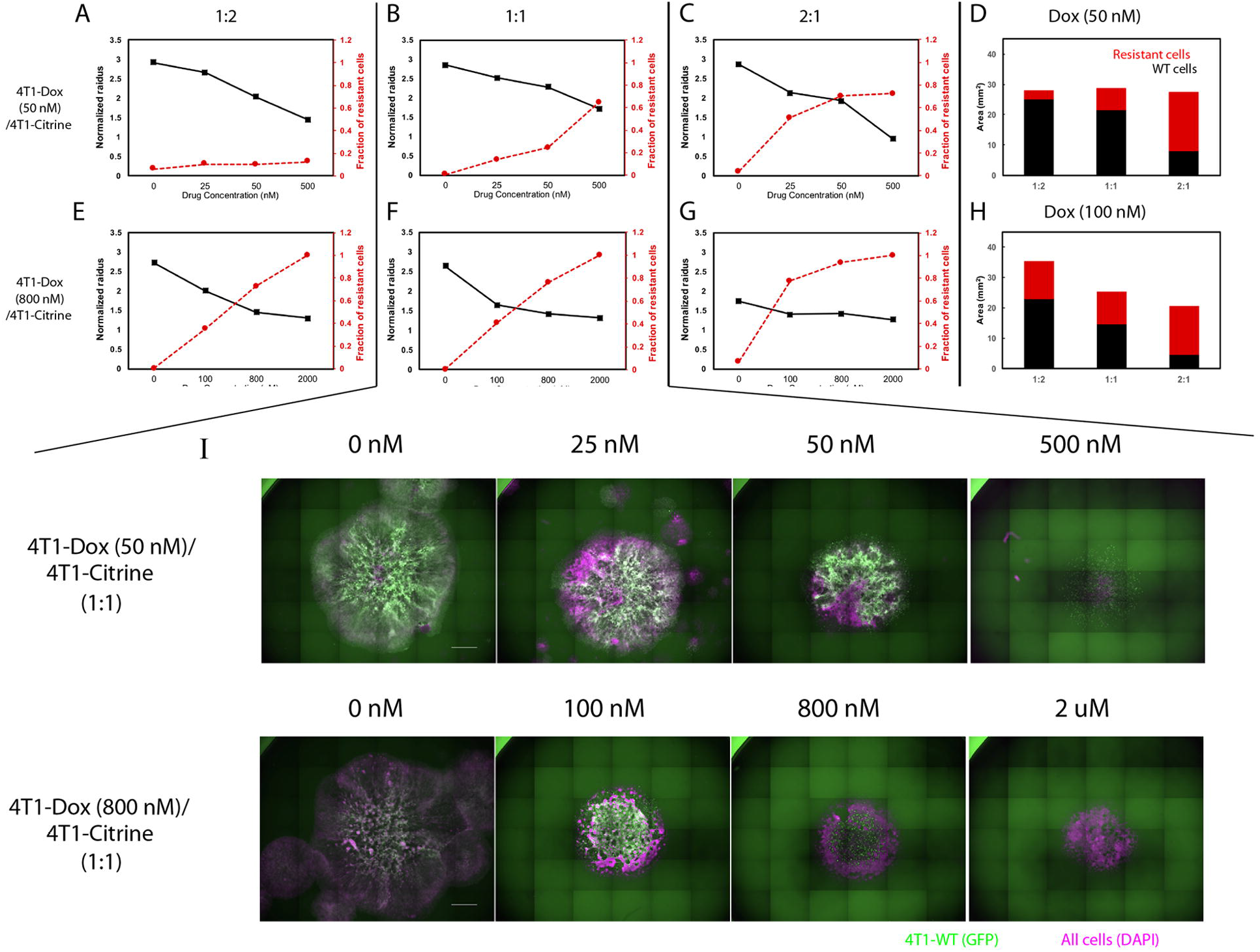
Relationship between overall 2D tumor colony size and treatment efficacy. **(A-C)** Normalized radius (black) and Fraction of resistant cells (red) for mixtures of chemosensitive 4T1 and 50 nM 4T1-Dox resistant cells at varied ratios. **(D)** Surface area of colony occupied by resistant (red) and chemosensitive cells (black). **(E-G)** Normalized radius (black) and Fraction of resistant cells (red) for mixtures of chemosensitive 4T1 and 800 nM 4T1-Dox resistant cells at varied ratios. **(H)** Surface area of colony occupied by resistant (red) and chemosensitive cells (black). **(I)** Stitched 4X fluorescent images of colonies demonstrating the distribution of chemosensitive cells (4T1-WT, GFP) in each colony (DAPI) in the presence of doxorubicin. Colonies were formed by seeding 1:1 mixture of 4T1-WT and 50 nM resistant cells (top, corresponding to B) or 1:1 mixture of 4T1-WT and 800 nM resistant cells (bottom, corresponding to F). Scale bar = 1000 μm.

As the size of the colony could not predict the efficacy of the sensitive and the resistant cells to the optimum resolution for a given dose of drug, we investigated if the schedule of drug treatment may alter the treatment efficacy. In order to study the effect of multi-dosage schedule on colony expansion, we seeded 4T1-WT cells and treated the colony with 3 different chemotherapy dosing schedules. Specifically, we varied the drug concentration and the frequency of treatment per day: 1 dose of 600 nM, 2 doses of 300nM or 3 doses of 200nM of doxorubicin. In this way, a cumulative treatment of 600 nM per day was achieved in each scenario. Gentle PBS wash was done for several times after each dose to remove waste in culture and fresh media with drug was then added for next dose. We found that when holding the cumulative treatment constant, colony expansion kinetics were not affected by altered dosing schedule or altered serum level in the media (Fig 4). Hence, the cumulative treatment but not the dose level determined the treatment efficacy in a serum-independent manner. Taken together, our model system can be utilized to identify a lower but effective drug concentration and can be utilized to test cumulative treatment regimes.

**Fig 4.**
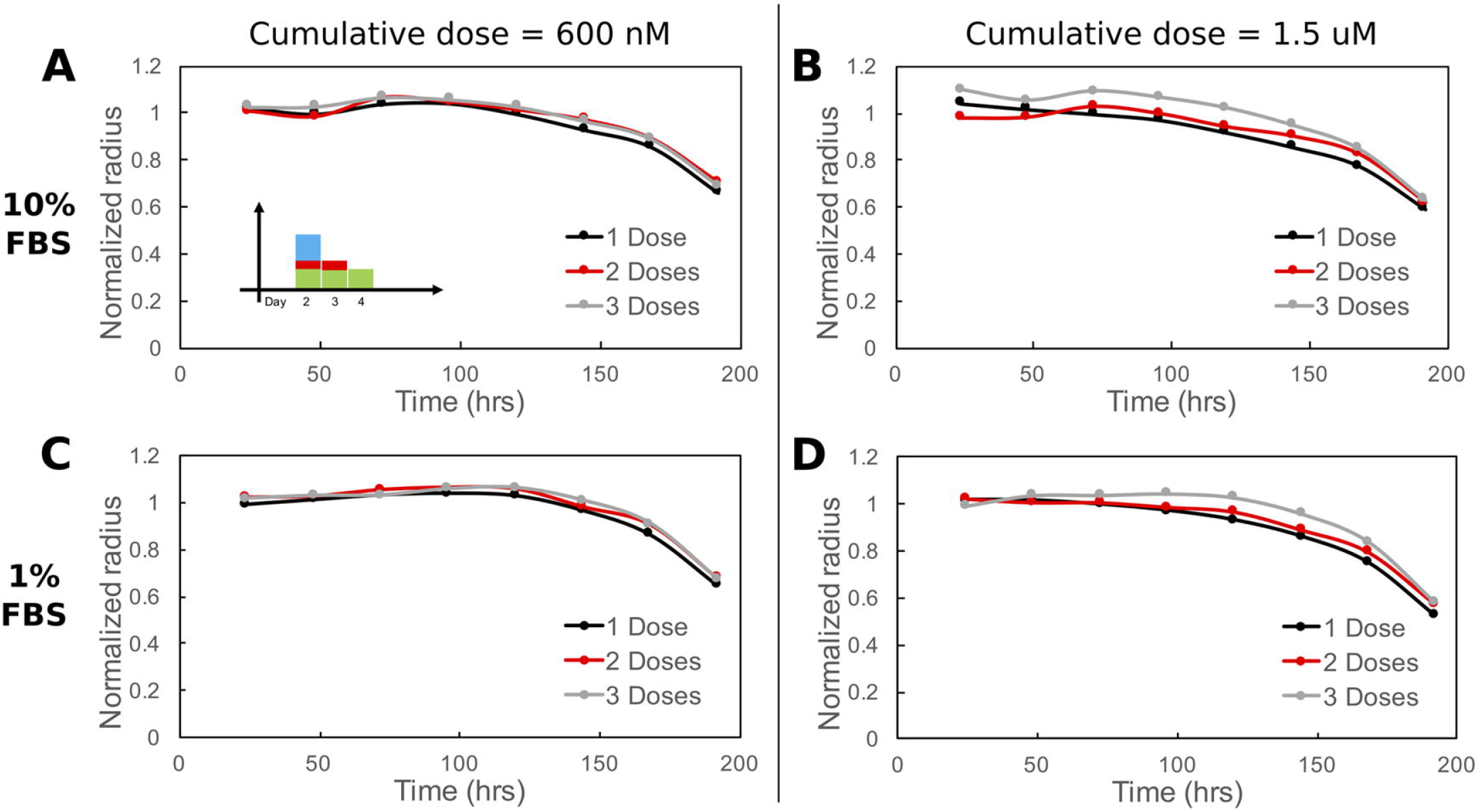
Cumulative treatment regime and long-term overall efficacy. **(A, C)** Normalized radius for 3 chemotherapeutic schedules of similar cumulative dose (600 nM) in 10% and 1% FBS. **(B, D)** Normalized radius for the same 3 chemotherapeutic schedules of similar cumulative dose (1.5 μM) in 10% and 1% FBS. Cumulative dosage for each chemotherapeutic schedule is defined as dosage level (nM) multiplied by duration of drug treatment (dosing frequency [1/day] × n days, n = 1, 2 or 3).

### A mathematical model based on the behavior and parameters of the 2D co-culture system can be used to predict cumulative chemotherapy regimens to attain the lowest volume of tumor

To quantitatively understand the dynamics of chemosensitive and chemoresistant cells subject to different dosing strategies, we built a mathematical model. In this model, chemosensitive (N_s_) and chemoresistant (N_r_) cells are initially mixed at a given ratio and then assumed to exponentially grow at rates μ_s_ and μ_r_, respectively (Fig 5A). To subject cells to chemotherapy, we used our experimentally determined doxorubicin dose-response viability curve for wildtype and resistant 4T1 cells (Fig 2A). Thus, cells after a given interval of time would have the population level N = V(x) * e^a^, where *V(x)* is the viability value for cells at a particular dose x, and *a* is the growth rate (a scaled time interval of t=1 is assumed here for simplicity). In the case of multiple doses, each dose is fractionated across a smaller interval, but repeated for multiple intervals, keeping the same total time interval as the single dose (Supplementary Information).

**Fig 5.**
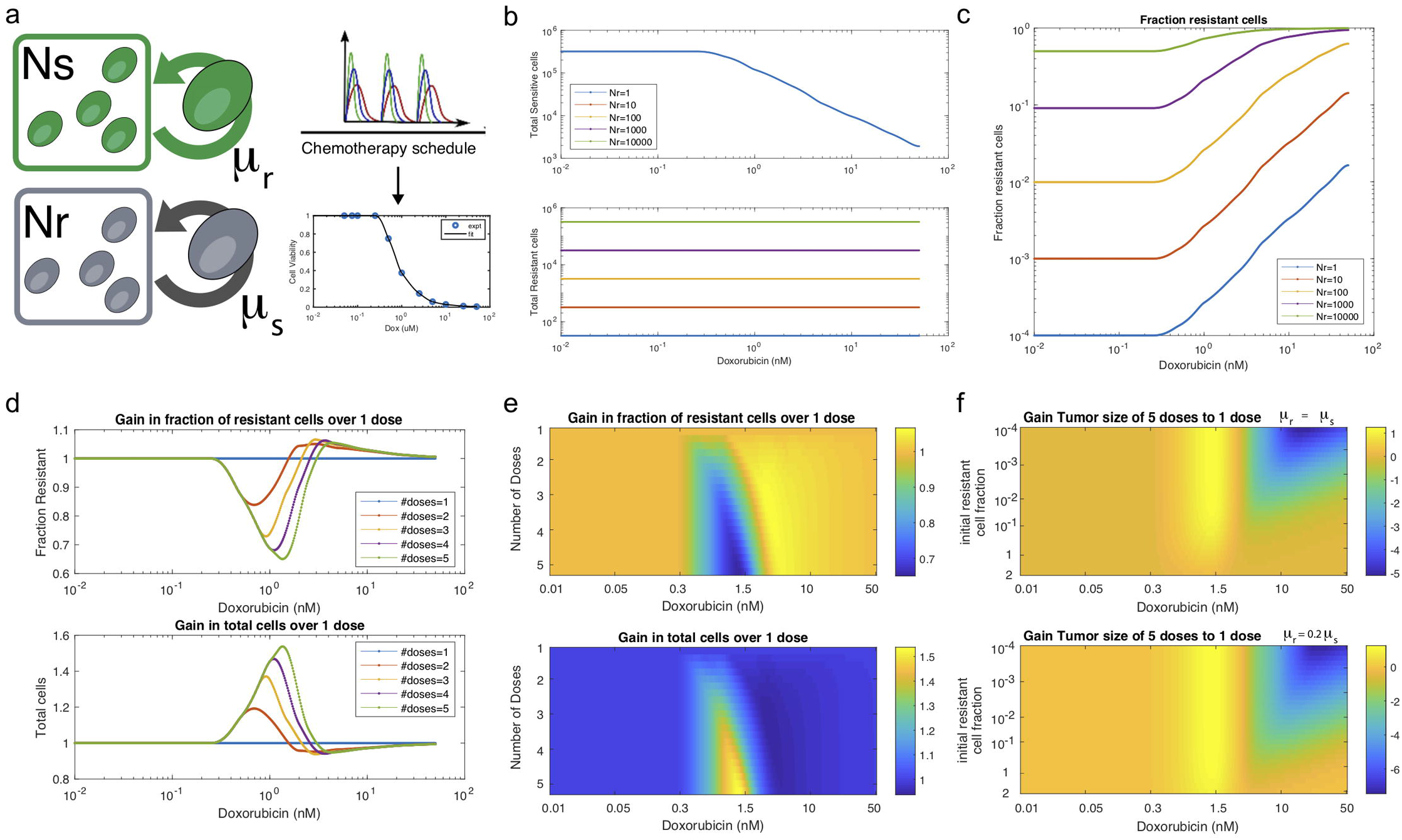
Mathematical model chemosensitive and chemoresistant populations. **(a)** Schematic of the underlying system modeled, whereby resistant and sensitive cell populations grow exponentially, and are then subject to chemotherapy regimens of varying frequency and dosage. **(b)** Total sensitive cells and total resistant cells at the end of interval as a function of starting dosage. Colors indicate varied initial conditions of Nr = 10^2^, 10^3^, 5×10^3^, 10^4^, 2×10^4^, where N_s_ = 10^4^. **(c)** Fraction of resistant cells as a function of starting dosage for varied initial conditions. **(d)** For a 1:1 mixture of N_s_ to N_r_, number of doses is varied. The fraction resistant, total cell population are plotted on the top row. The ratio of a particular number of doses to 1 dose is termed the “Gain”, both for fraction resistant and total cell population. **(e)** Gains are given as a function of starting dose (x axis) and number of doses (y axis).

We first simulated the model and varied the number of resistant cells N_r_ from 10^3^-2×10^4^, while keeping the number of chemosensitive cells at 10^4^. Here we observed that the total number of chemosensitive cells was reduced as we increased the initial dosage for a single dose, as expected, while the amount of chemoresistant cells was not affected by chemotherapy dose (Fig. 5B). This latter result is because we simulated highly resistant cells such as 4T1 (Dox 3uM) are viable at all chemotherapy doses tested. We then calculated the fraction of resistant cells across multiple doses and initial conditions (Fig. 5C). For high doses, such as in a maximum tolerated dose (MTD) approach, resistant populations eventually dominate the cell population as previously noted. Next, we set sensitive and resistant cells at a 1:1 initial ratio and assumed similar growth rates. We found that when varying the number of doses applied, the fraction of resistant cells and total cell number as compared to a single dose (termed “gain”) was a function of the initial starting dose of chemotherapy used (Fig. 5D). Low concentrations of chemotherapy resulted in populations with a lower fraction of resistant cells when comparing multiple doses to a single dose (low gain). In turn, multiple doses also resulted in larger total cell populations (high gain), which can be interpreted as larger tumor sizes. However, at high concentrations of chemotherapy, the relationship was observed to be reversed. Higher levels of chemotherapy with multiple doses led to an increased fraction of resistant cells and smaller total cell number. The relationship between increasing resistant cell fraction and smaller total cell number was inversely proportional, such that there was no particular number of doses or starting dose that would allow for beneficial optimization of both measures (Fig 5E). Lastly, dependent on the initial resistant cell fraction, we found that sharper gains could be obtained with less numbers of initial resistant cells. This trend was also further magnified when resistant cells grew slower (Fig 5F). A common clinical situation is when initial resistant cell populations are low and slow growing, and in these cases the effect of metronomic dosing may be magnified. In summary, based on the kinetics of our 2D model system, this simple mathematical model can predict a critical concentration of drug and a schedule of treatment to achieve the smallest volume of tumor, the lowest proportion of resistant cells, or in between points that balance the trade-off between these outcomes.

### The 2D co-culture system with chemosensitive and chemoresistant cells can be optimized to create spatial heterogeneity found in 3D tumors

Here, we tested if growth of chemo-resistant cell populations can be affected by the spatial organization of chemo-sensitive cells in our co-culture system. We observed that the chemoresistant cell’s growth was constrained by surrounding chemosensitive cells and the constrained growth was largely depended on their inoculated location (Fig 6, Fig S3-S7). When resistant cells located in the center of the colony, surrounding sensitive cells outcompeted resistant cells for space and prevented resistant colony expansion. In contrast, a semi-constrained scenario was that when resistant cells located near the edge of the colony, only inward invasion of resistant colony was prohibited by surrounding sensitive cells while outward expansion was still possible. This drastic difference between the complete and semi-constrained scenarios was more obvious when two resistant cell subpopulations were inoculated in the same colony (Fig S7). Hence, our 2D co-culture system can be optimized to imitate various tumor architectures with different layers of spatial heterogeneity.

**Fig 6.**
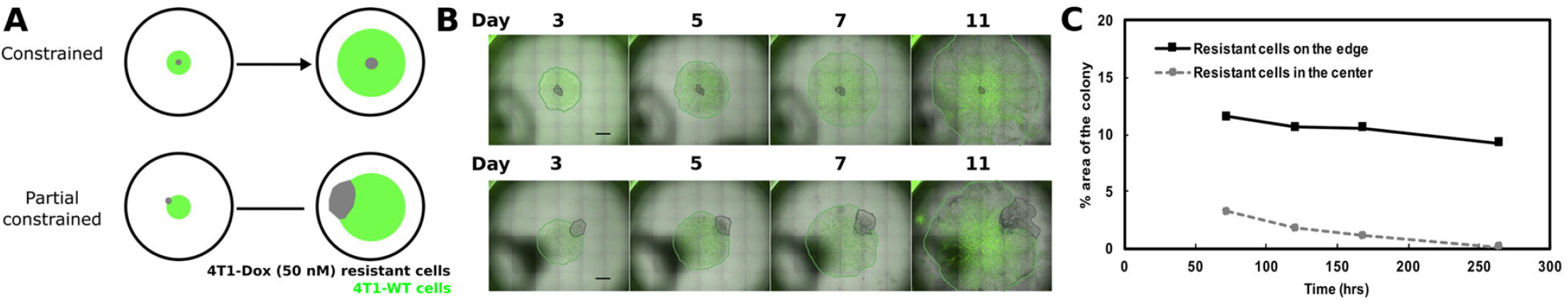
Spatial heterogeneity in 2D co-culture system created by varying the initial inoculation location. **(A)** Schematic demonstrating how placement of the resistant population can affect colony structure formation over time. **(B)** 20X fluorescent and brightfield composites of 2D colony of chemosensitive and resistant cells. **(C)** Fraction of resistant cells over time on the edge and center of the colony.

## Discussion

Several clinical trials in the past have shown that the maximum tolerated dose approach has not always provide maximum clinical benefits. For example, Cisplatin, a common chemotherapy administered in non-small cell lung cancer patients, failed to show any clinical benefit in terms of overall survival or pathological complete response over relatively moderate doses in a randomized multicenter trial (27). Furthermore, retrospective analyses on low dose metronomic chemotherapy, with frequent schedules proved to be clinically favorable and safer compared to conventional chemotherapy for a large number of drugs in a broad range of tumors (28). In our 2D co-culture system of doxorubicin-sensitive and resistant 4T1 cells, we observed that a cumulative treatment regimen with low dose administered in frequent interval resulted in the same size of colonies as in the regimen with high dose administered altogether (Fig 4). As our simple 2D coculture model enabled to visually track the subpopulation of chemosensitive and chemoresistant cells for various treatment regimens over time, we found that despite showing same colony size, the growth dynamics of these sub-populations were radically different. Furthermore, we observed different behavior of chemoresistant cells based on their level of resistance. 4T1-Dox [800 nM], 4T1 cells that could survive 800 nM of doxorubicin, increased over time similar to 4T1 cells that survived 50 nM doxorubicin. But, once the number of the 4T1-wildtype and 4T1-Dox [800 nM] subpopulations reach the same number, increasing the dose of doxorubicin did not reduce the overall colony size markedly (Fig 3). Hence, our simple 2D co-culture system could be optimized to identify lowest effective dose based on the growth kinetics of chemosensitive and chemoresistant cells.

The challenges associated with experimentally tracking growth and response of chemosensitive and chemoresistant cells in a mixed culture has inspired mathematical modeling of tumor architectures (29–33). Recently, a study used an off-lattice agent-based model and flow-cytometry analysis to show the benefits of adaptive therapeutic strategies using dose modulation or treatment vacation in breast cancer (19). Compared to such complex simulations, we took a very simple approach of building our model based on the dose response curves of chemosensitive and chemoresistant cells from the *in vitro* co-culture. Nonetheless, we successfully identified a treatment regimen to attain the smallest sized tumor. The simplicity of our model also allows us to further optimize it for other cancer types and other chemotherapy and targeted therapy drugs quickly.

As our 2D co-culture system enables easy tracking of the growth dynamics of chemosensitive and chemoresistant cells under simple epifluorescence microscopy, we can further optimize them for high throughput screening purposes to identify optimal combinatorial therapies and discovery of new treatment strategies via screening of experimental small molecule libraries (34). Our 2D co-culture system also provides us the control to build various types of spatial architecture by simply changing the location of initial inoculation (Fig 6). Different architectures quickly created by this model can easily represent complex patterns of spatial heterogeneity that may vary from one patient to another. Together, with mathematical modeling, our simple system to build 2D cocultures provides an effective approach to find optimum therapeutic regimen for heterogenous tumors.

## Acknowledgement

This study was supported by NIH Pathway to Independence Award (R00CA197649-02) and DoD Era of Hope Scholar Award (BC160541).

## Supporting Information

**Fig S1 Radial growth dynamics of 4T1-WT monoculture**. Colony radius as a function of time for various seeding densities in the presence of **(A)** 10%, **(C)** 5%, or **(E)** 1% FBS. Growth dynamic data were fitted using linear regression, slopes and intercepts of regression lines (dotted) are listed in **(B)** 10% FBS, **(D)** 5% FBS, and **(F)** 1% FBS.

**Fig S2. Growth dynamics of mixed binary colonies in the presence doxorubicin**. Colony radius normalized to starting radius in a 1:2 **(A)**, 1:1 **(B)** or 2:1 **(C)** seeded mixture of sensitive and 50nM Dox resistant cells in the presence of 0 nM, 25 nM, 50 nM or 500 nM doxorubicin. Colony radius normalized to starting radius in a 1:2 **(D)**, 1:1 **(E)** or 2:1 **(F)** seeded mixture of sensitive and 800nM Dox resistant cells in the presence of 0 nM, 100 nM, 800 nM or 2 μM doxorubicin.

**Fig S3. Modulation of spatial competition in colonies where resistant cells were located in the center. (A)** Stitched 4X fluorescent and bright field composites demonstrating the distribution of chemosensitive cells (4T1-WT, GFP) and 50 nM doxorubicin resistant cells (grey) over time in the presence of 10 % FBS. Colonies were formed by seeding 9:1 of 4T1-WT and chemoresistant cells, chemoresistant cells were seeded in the center of the colony. Colonies were either untreated (top) or treated with two 500 nM doxorubicin doses at day 5 and day 9 (bottom). **(B)** % area of resistant cells over time in control or doxorubicin treated colonies. Scale bar = 1000 μm.

**Fig S4. Modulation of spatial competition in colonies where resistant cells were located in the center. (A)** Stitched 4X fluorescent and bright field composites demonstrating the distribution of chemosensitive cells (4T1-WT, GFP) and 50 nM doxorubicin resistant cells (grey) over time in the presence of 1% FBS. Colonies were formed by seeding 9:1 of 4T1-WT and chemoresistant cells, chemoresistant cell were seeded in the center of the colony. Colonies were either untreated (top) or treated with two 500 nM doxorubicin doses at day 5 and day 9 (bottom). **(B)** % area of resistant cells over time in control or doxorubicin treated colonies. Scale bar = 1000 μm.

**Fig S5. Modulation of spatial competition in colonies where resistant cells were located on the edge (A)** Stitched 4X fluorescent and bright field composites demonstrating the distribution of chemosensitive cells (4T1-WT, GFP) and 50 nM doxorubicin resistant cells (grey) over time in the presence of 10% FBS. Colonies were formed by seeding 9:1 of 4T1-WT and chemoresistant cells, chemoresistant cell were seeded on the edge of the colony. Colonies were either untreated (top) or treated with two 500 nM doxorubicin doses at day 5 and day 9 (bottom). **(B)** % area of resistant cells over time in control or doxorubicin treated colonies. Scale bar = 1000 μm.

**Fig S6. Modulation of spatial competition in colonies where resistant cells were located on the edge (A)** Stitched 4X fluorescent and bright field composites demonstrating the distribution of chemosensitive cells (4T1-WT, GFP) and 50 nM doxorubicin resistant cells (grey) over time in the presence of 1% FBS. Colonies were formed by seeding 9:1 of 4T1-WT and chemoresistant cells, chemoresistant cell were seeded on the edge of the colony. Colonies were either untreated (top) or treated with two 500 nM doxorubicin doses at day 5 and day 9 (bottom). **(B)** % area of resistant cells over time in control or doxorubicin treated colonies. Scale bar = 1000 μm.

**Fig S7. Modulation of spatial competition in colonies. (A)** Stitched 4X fluorescent and bright field composites demonstrating the distribution of chemosensitive cells (4T1- WT, GFP) and 50 nM doxorubicin resistant cells (grey) over time in the presence of 10% FBS. Colonies were formed by seeding 8:2 of 4T1-WT and chemoresistant cells, half of chemoresistant cells were seeded in the center while the other half were seeded on the edge of the colony. Colonies were either untreated (top) or treated with two 500 nM doxorubicin doses at day 5 and day 9 (bottom). **(B)** % area of resistant cells over time in control or doxorubicin treated colonies. Scale bar = 1000 μm.

