## Supporting figures, legends and text for "A spatial cell culture model for predicting chemotherapy dosing strategies"

**Figure S1**

**A**

**10% FBS**

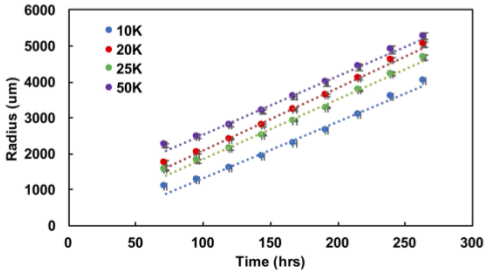


**D**

| Seeded cells | Slope | Intercept | $R^2$ |
|---|---|---|---|
| 10K | 15.55 | -223.85 | 0.99 |
| 20K | 17.43 | 356.56 | 0.99 |
| 25K | 16.53 | 206.61 | 0.99 |
| 50K | 16.28 | 915.48 | 0.99 |

**B**

**5% FBS**

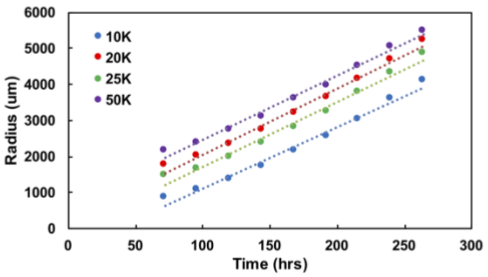


**E**

| Seeded cells | Slope | Intercept | $R^2$ |
|---|---|---|---|
| 10K | 17.12 | -596.61 | 0.98 |
| 20K | 18.37 | 223.25 | 0.99 |
| 25K | 18.06 | -83.01 | 0.98 |
| 50K | 17.879 | 666.14 | 0.99 |

**C**

**1% FBS**

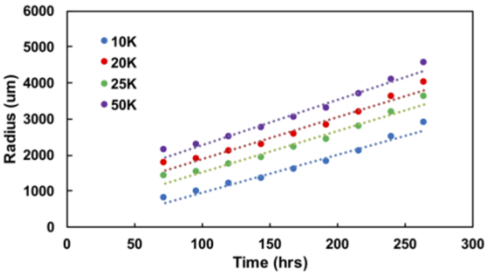


**F**

| Seeded cells | Slope | Intercept | $R^2$ |
|---|---|---|---|
| 10K | 10.56 | -91.09 | 0.97 |
| 20K | 11.81 | 704.54 | 0.97 |
| 25K | 11.42 | 389.65 | 0.97 |
| 50K | 12.52 | 1031.2 | 0.98 |

**Fig S1. Radial growth dynamics of 4T1-WT monoculture.** Colony radius as a function of time for various seeding densities in the presence of **(A)** 10%, **(C)** 5%, or **(E)** 1% FBS. Growth dynamic data were fitted using linear regression, slopes and intercepts of regression lines (dotted) are listed in **(B)** 10% FBS, **(D)** 5% FBS, and **(F)** 1% FBS.

### Figure S2

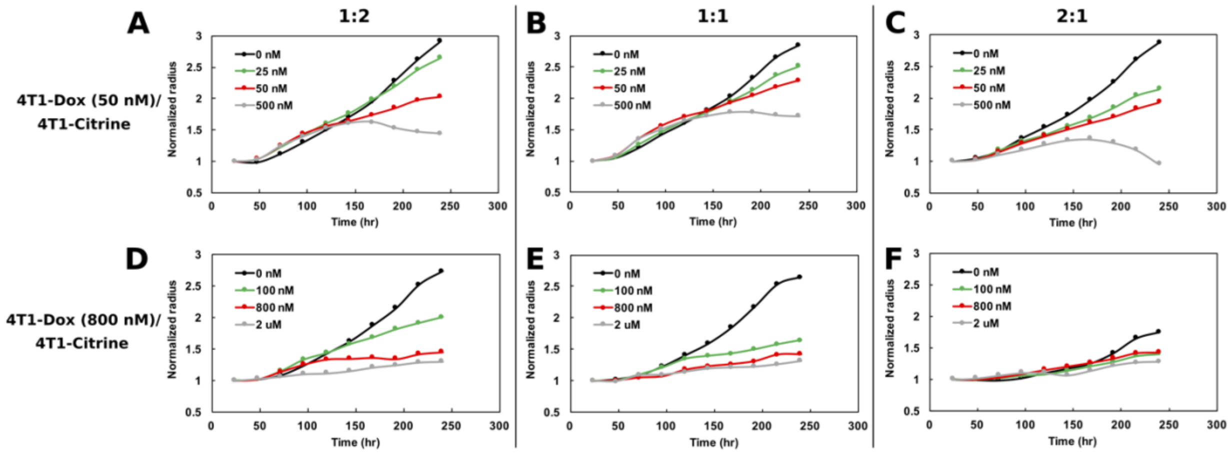


**Fig S2. Growth dynamics of mixed binary colonies in the presence doxorubicin.** Colony radius normalized to starting radius in a 1:2 **(A)**, 1:1 **(B)** or 2:1 **(C)** seeded mixture of sensitive and 50nM Dox resistant cells in the presence of 0 nM, 25 nM, 50 nM or 500 nM doxorubicin. Colony radius normalized to starting radius in a 1:2 **(D)**, 1:1 **(E)** or 2:1 **(F)** seeded mixture of sensitive and 800nM Dox resistant cells in the presence of 0 nM, 100 nM, 800 nM or 2 uM doxorubicin.

**Figure S3**

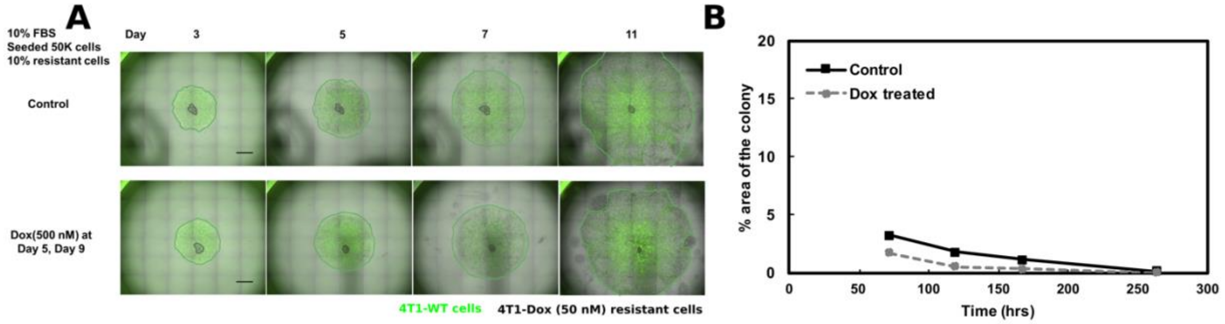


**Fig S3. Modulation of spatial competition in colonies where resistant cells were located in the center. (A)** Stitched 4X flourescent and bright field composites demonstrating the distribution of chemosensitive cells (4T1-WT, GFP) and 50 nM doxorubicin resistant cells (grey) over time in the presence of 10 % FBS. Colonies were formed by seeding 9:1 of 4T1-WT and chemoresistant cells, chemoresistant cells were seeded in the center of the colony. Colonies were either untreated (top) or treated with two 500 nM doxorubicin doses at day 5 and day 9 (bottom). **(B)** % area of resistant cells over time in control or doxorubicin treated colonies. Scale bar = 1000 um.

**Figure S4**

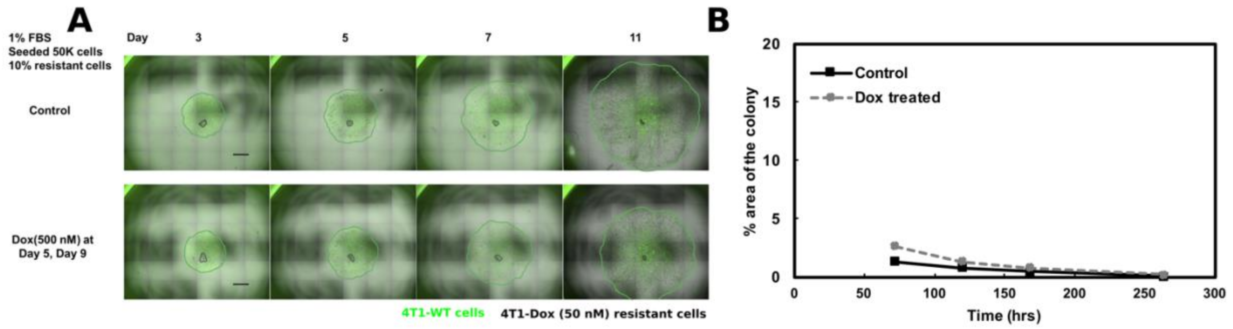


**Fig S4. Modulation of spatial competition in colonies where resistant cells were located in the center. (A)** Stitched 4X flourescent and bright field composites demonstrating the distribution of chemosenstive cells (4T1-WT, GFP) and 50 nM doxorubicin resistant cells (grey) over time in the presence of 1% FBS. Colonies were formed by seeding 9:1 of 4T1-WT and chemoresistant cells, chemoresistant cell were seeded in the center of the colony. Colonies were either untreated (top) or treated with two 500 nM doxorubicin doses at day 5 and day 9 (bottom). **(B)** % area of resistant cells over time in control or doxorubicin treated colonies. Scale bar = 1000 um.

**Figure S5**

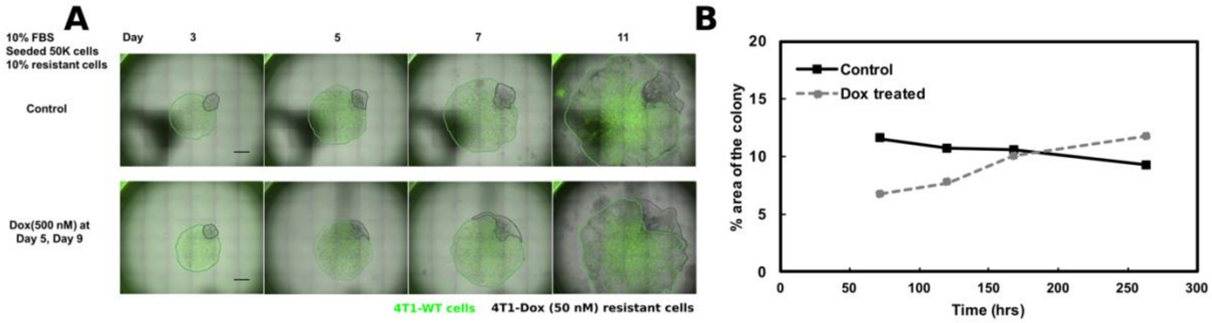


**Fig S5. Modulation of spatial competition in colonies where resistant cells were located on the edge (A)** Stitched 4X flourescent and bright field composites demonstrating the distribution of chemosenstive cells (4T1-WT, GFP) and 50 nM doxorubicin resistant cells (grey) over time in the presence of 10% FBS. Colonies were formed by seeding 9:1 of 4T1-WT and chemoresistant cells, chemoresistant cell were seeded on the edge of the colony. Colonies were either untreated (top) or treated with two 500 nM doxorubicin doses at day 5 and day 9 (bottom). **(B)** % area of resistant cells over time in control or doxorubicin treated colonies. Scale bar = 1000 um.

**Figure S6**

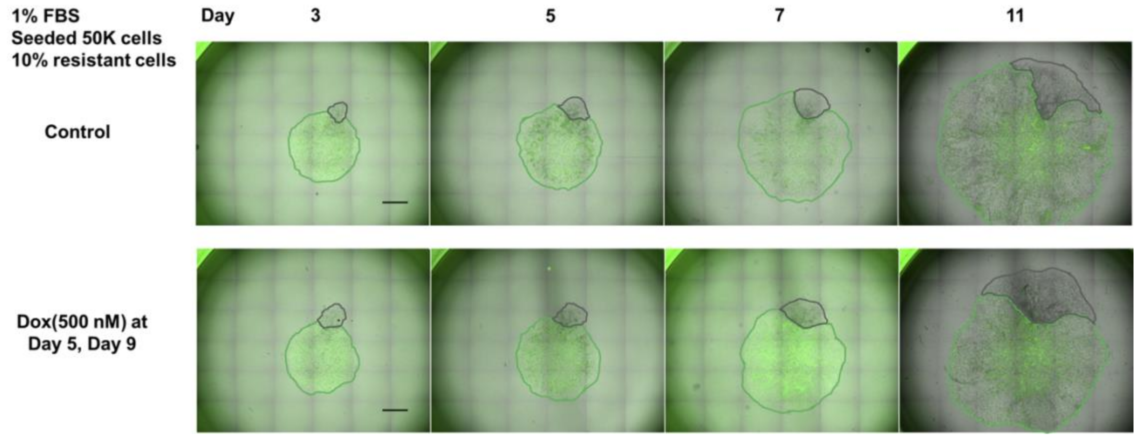


**Fig S6. Modulation of spatial competition in colonies where resistant cells were located on the edge (A)** Stitched 4X flourescent and bright field composites demonstrating the distribution of chemosenstive cells (4T1-WT, GFP) and 50 nM doxorubicin resistant cells (grey) over time in the presence of 1% FBS. Colonies were formed by seeding 9:1 of 4T1-WT and chemoresistant cells, chemoresistant cell were seeded on the edge of the colony. Colonies were either untreated (top) or treated with two 500 nM doxorubicin doses at day 5 and day 9 (bottom). **(B)** % area of resistant cells over time in control or doxorubicin treated colonies. Scale bar = 1000 um.

**Figure S7**

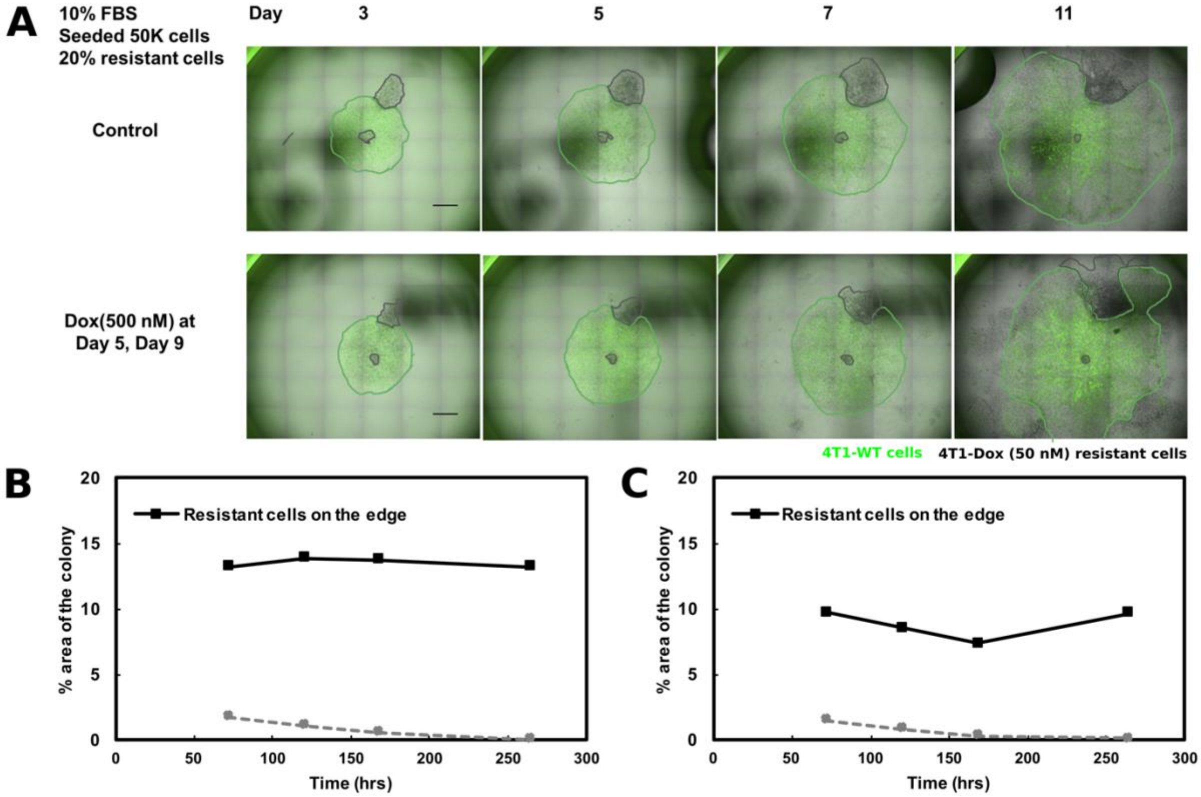


**Fig S7. Modulation of spatial competition in colonies. (A)** Stitched 4X flourescent and bright field composites demonstrating the distribution of chemosenstive cells (4T1-WT, GFP) and 50 nM doxorubicin resistant cells (grey) over time in the presence of 10% FBS. Colonies were formed by seeding 8:2 of 4T1-WT and chemoresistant cells, half of chemoresistant cells were seeded in the center while the other half were seeded on the edge of the colony. Colonies were either untreated (top) or treated with two 500 nM doxorubicin doses at day 5 and day 9 (bottom). **(B)** % area of resistant cells over time in control or doxorubicin treated colonies. Scale bar = 1000 um.

**Mathematical model of chemosensitive and chemoresistant cells**

The analytical model for chemosensitive and chemoresistant cell population growth is described in the main text. The model assumes exponential growth of both populations with no competing interactions or limited nutrients. In this simplified model, for each starting dose (x), cells were assumed to change in number given by the fraction of cells viable given in the dose-response curve V(x). For instance, at 2 μM of doxorubicin (Fig 2A), V(2)=0.5, and thus ~50% of WT cells survived for each dose applied. More generally, the number of cells after a single dose of chemotherapy is given by the equation:

$N = V(x)*e^a$ (1)

where *a* is the growth rate and a scaled time interval of t=1 is assumed here for simplicity. In the case of multiple doses, each dose is fractionated across the interval, and the dose is given by X/n, where *n* is the number of doses. In comparison to a single dose, the total number of cells for multiple doses would be

$N = [V(X/n)*e^{a/n}]^n$ (2)

This is also equivalent to

$N= V(X/n)^n*e^a$ (3)

Comparing equation 3 with equation 1 shows that the ratio of the number of cells after multiple doses compared to a single dose (termed the gain) is solely dependent on the dose-response curve values and not on growth rate of cells. Here, if $V(x) < V(x/n)^n$, then a single dose will produce less cells (and hence a smaller tumor) than multiple doses. We applied this equation to both chemosensitive and chemoresistant populations by using the experimentally determined Doxorubicin dose-reponse curves in Fig 2A (WT and 4T1 – 3uM).
